# VelocityFM: Short-Horizon Protein Trajectory Prediction via Flow Matching in Velocity Space

**DOI:** 10.64898/2026.06.05.730410

**Authors:** G.D.L.U. Jayathilake, C. R. Wijesinghe, A. R Weerasinghe

## Abstract

Protein dynamics is fundamentally a trajectory prediction problem, but molecular dynamics (MD) simulation remains expensive and static structure predictors do not model time-ordered motion. We present *VelocityFM*, a short-horizon protein trajectory predictor that applies rectified flow matching in velocity space over residue frames and torsions. The model combines six Invariant Point Attention (IPA) blocks with a two-layer per-residue temporal self-attention encoder, and is trained on 710 ATLAS proteins comprising 2090 filtered replicate trajectories. At the primary 128-frame rollout horizon, VelocityFM achieves a median TM-score of 0.929 on 72 held-out proteins, with 100% of proteins remaining above TM> 0.7 and 100% clash-free generation. Backbone geometry also remains strong, with a median Ramachandran favoured rate of 91.09%, while dynamics calibration is conservative with median RMSF ratio 0.697. These results show that velocity-space geometric learning can generalise short-horizon trajectory prediction to unseen proteins while preserving fold structure and geometric validity within its intended operating regime.

## 1 Introduction

Proteins function through motion as much as through structure. Catalysis, allostery, gating, and binding all depend on transitions between nearby conformational states rather than on a single static fold [1]. Molecular dynamics (MD) simulation provides physically grounded trajectories, but its cost scales poorly with the timescales of interest, creating a practical gap between structural prediction and dynamical modelling [2, 3]. From a machine learning viewpoint, this motivates short-horizon predictors that can learn local trajectory evolution directly from MD data instead of reproducing full long-horizon simulation.

A useful learned dynamics model must satisfy several constraints at once. It should preserve global fold, maintain local geometry, avoid steric failure, and produce motion that is temporally ordered rather than merely a set of diverse conformations. This makes the problem different from standard single-structure prediction and also different from ensemble generation. The target object is a short stochastic process over 3D states.

We study this problem with *VelocityFM*, a residue-level generative model for short-horizon trajectory prediction. Instead of predicting Cartesian coordinates directly, VelocityFM predicts translational, rotational, and torsional *velocities* on residue frames, then integrates those updates on SO(3) and on wrapped torsion angles. This formulation is attractive for dynamics because it makes the prediction target local in time, reduces target variance relative to absolute coordinate prediction, and matches the way trajectories are numerically propagated.

The model uses six IPA blocks for geometry-aware spatial reasoning and a two-layer per-residue temporal attention encoder over windows of *W* =16 frames. It is trained with rectified flow matching in velocity space on 710 proteins and 2090 filtered ATLAS replicate trajectories, using a staged loss curriculum that delays heavy geometry losses until the velocity field is stable. On the held-out 72-protein test set at the primary 128-frame rollout horizon, the median TM-score is 0.929, all proteins remain above TM*>* 0.7, and all generated rollouts are clash-free. The same evaluation yields a median Ramachandran favoured rate of 91.09% and a median RMSF ratio of 0.697, showing strong structural preservation with conservative motion amplitude.

These results suggest a specific interpretation. VelocityFM is not a replacement for MD, and it is not yet a long-horizon simulator. It is instead a strong short-horizon predictor that generalises fold-preserving local trajectory evolution to unseen proteins. The goal of this paper is to describe the architecture and training design that make this possible, and to characterise clearly where the model succeeds and where it remains limited.

### Contributions

Our main contributions are:

- a velocity-space flow matching formulation for short-horizon protein trajectory prediction over residue frames and torsions,
- an IPA + temporal attention architecture that couples geometry-aware spatial reasoning with per-residue temporal modelling,
- a staged loss curriculum with automatic *σ*^2^ normalisation for physical-space losses, improving optimisation stability under mixed supervision, and
- a 72-protein held-out evaluation demonstrating complete fold recognition generalisation at 128 frames, with 100% TM > 0.7 and 100% clash-free generation.

## 2 Related Work

MD simulation remains the reference method for protein dynamics because it explicitly integrates time-evolving atomic motion, but its cost limits routine access to longer trajectories and larger systems [2, 3]. Learned trajectory and ensemble models try to approximate parts of this behaviour directly from data. Recent examples include MDGen, which models MD trajectories with a generative sequence model, and AlphaFlow, which uses flow matching to generate conformational ensembles rather than ordered rollouts [4, 5]. In our setting, the key distinction is that the target object is a short, time-ordered trajectory conditioned on a starting state.

Protein generative modelling has also moved strongly toward SE(3)-aware representations. AlphaFold established frame-based geometric reasoning at scale, and AlphaFold 3 extended the paradigm to broader biomolecular structure prediction [6, 7]. RFdiffusion and Boltz-1 show that equivariant generative models can be effective for structured 3D biomolecular outputs, but they target structure generation or interaction modelling rather than MD-style temporal prediction [8, 9]. VelocityFM adopts the same geometric ML viewpoint while making time-ordered state evolution the central objective.

Flow matching provides a practical continuous-time training objective for generative modelling by regressing vector fields along interpolants between noise and data [10]. In proteins, AlphaFlow, P2DFlow, and related work show that flow-based objectives can model conformational distributions and protein ensembles [5, 11]. Our approach differs in two ways: it learns in *velocity space* rather than in structure space, and it combines frame-aware spatial attention with explicit temporal attention over windows of residue trajectories.

## 3 Method

### 3.1 Problem Formulation

We represent a protein with *N* residues at time *t* as

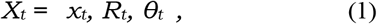

here *x*_*t*_ ∈ R^*N*×3^ stores residue translations, *R*_*t*_ ∈ SO(3)^*N*^ stores residue-frame orientations, and *θ*_*t*_ ∈ R^*N*×7^ stores seven torsion angles per residue. Given a conditioning state 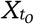, the model predicts a future window

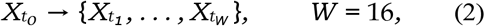

using multi-stride supervision with strides {1, 2, 4}.

The supervised targets are not absolute future states but per-step velocities:

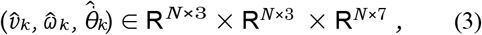

corresponding to translational velocity, body-frame angular velocity, and torsion velocity. With step size Δ*t*, the targets are computed from consecutive states as

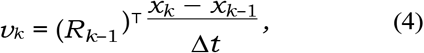

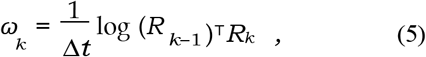

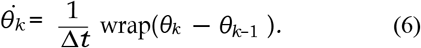

This representation makes the learning target local in time, expresses motion in the residue frame rather than in an arbitrary global coordinate system, and decouples global pose from short-horizon state change.

In the final implementation, the frame origin is taken from the residue C*α* position and the orientation is derived from backbone geometry. This gives every residue a compact rigid-body state while leaving fine local flexibility to the torsion variables. The decomposition is important because it separates two different sources of motion: larger coordinated displacement captured by (*x, R*) and finer internal change captured by *θ*.

### 3.2 Flow Matching in Velocity Space

For each training window we form a target velocity tensor *y*_1_ from MD-derived translational, rotational, and torsional velocities, and a noise sample *y*_0_ from an AR(1)-correlated Gaussian prior along the temporal dimension. If *ρ* denotes the temporal correlation coefficient and *ε*_*k*_ ∼ N (0, *I*), the base process can be written as

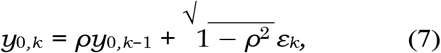

with component-specific scales for translational, rotational, and torsional channels.

Rectified flow matching uses the linear interpolant

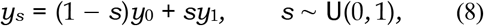

and trains a network *f*_*θ*_ to regress the conditional vector field. The core objective is

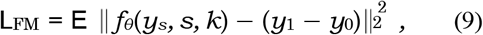

where *k* denotes the step index within the window. In practice, all velocity channels are normalised before regression, so the FM loss is already an *O*(1) objective even though the physical channels have different units.

Velocity space is preferable to direct coordinate-space prediction here for two reasons. First, the target is lower variance because neighbouring MD frames are close in time even when their absolute poses vary across proteins. Second, residue-frame velocities match the downstream integration step used during inference, so the training target and the rollout mechanism operate in the same representation. The model therefore learns how state changes, not merely where future coordinates happen to lie.

A practical consequence is that the same flow formulation can be used across proteins of very different sizes and structural classes without asking the network to memorise absolute coordinate templates. The regression target is a change field conditioned on the current state, which is a better match to short-horizon rollout than a direct next-frame coordinate map.

### 3.3 Architecture

VelocityFM instantiates the model class `VelocityFlowIPADynamics` with approximately 15M parameters. Each residue receives a 36-dimensional input vector formed by concatenating the conditioning state and the current flow state:

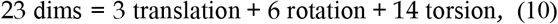

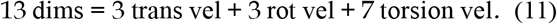

The rotation is represented with a 6D continuous embedding, and the torsions use sine/cosine encoding so that periodicity is respected before projection.

The 36-dimensional vector is projected to the single representation dimension *C*_*S*_ = 384 and augmented with flow time *s*, step index *k*, stride embedding, and a linear projection of frozen ESM2 residue embeddings from 1280 → 384 [12]. Pair features are stored in a representation of size *C*_*Z*_ = 128 and encode residue-residue geometric context used by the spatial trunk. In the final implementation, the pair tensor is shared across the temporal window so that the network does not recompute static residue-pair geometry at every step.

Spatial reasoning is handled by six IPA blocks with 8 attention heads, 4 query/key points, and 8 value points, with numerically sensitive internal computations kept in FP32 inside an otherwise BF16 run. The IPA stack operates on per-residue features and pair features, giving the model equivariant access to local and non-local geometry [6]. This is the main source of geometry awareness in the model: residues attend not only through feature similarity but also through their learned relation to 3D frame geometry. In practice, this lets the network couple sequence-conditioned residue identity with geometry-conditioned frame context.

After the spatial encoder, a two-layer per-residue self-attention module with 8 heads runs across the *W* = 16 temporal axis. The tensor is reshaped so that each residue attends only across its own trajectory history inside the window. This factorisation is important: the spatial encoder handles residue–residue coupling within a frame, whereas the temporal encoder handles frame–frame dependence for each residue across time. The temporal block therefore learns whether a residue is continuing a trend, reversing direction, or remaining approximately stationary across recent steps.

The model ends with stride-specific linear heads for strides {1, 2, 4}, each predicting translational, rotational, and torsional velocities. Separate heads avoid forcing a single output projection to represent multiple physical step sizes simultaneously. In effect, the model shares the trunk across timescales but allows the final mapping to remain stride-aware. This is especially useful because a stride-4 target should not be treated as a simple rescaling of a stride-1 target once rotations, torsions, and geometry losses are all present. The architecture of VelocityFM is shown in Figure 1.

**Figure 1:**
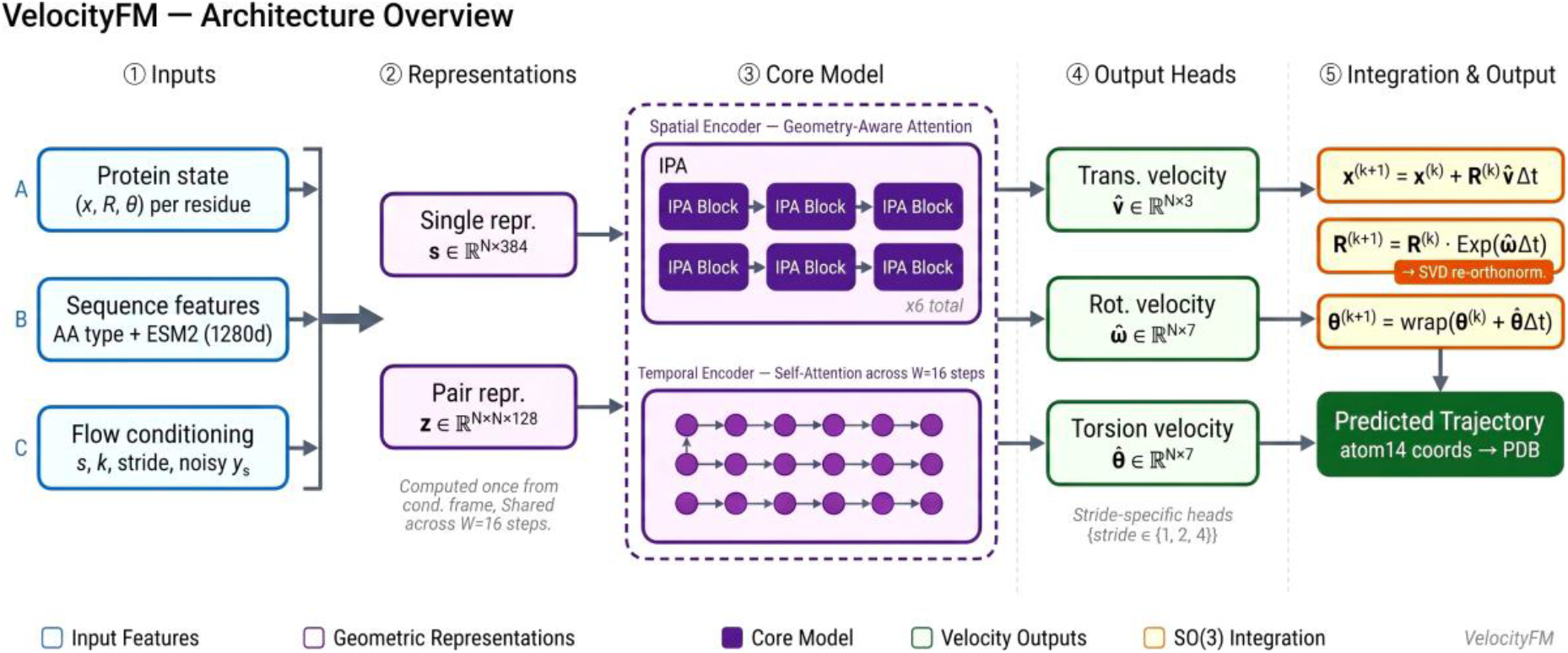
VelocityFM architecture. Input residue states are projected into a 36-dim per-residue vector, processed by six IPA blocks for geometry-aware spatial interaction, a temporal attention encoder across *W* =16 steps, and stride-specific velocity heads. Predicted velocities are integrated on SO(3) to produce output trajectory frames.

### 3.4 SO(3) Integration and Reconstruction

Predicted velocities are integrated with first-order Euler updates in physical time. For residue *i* at step *k*,

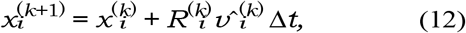

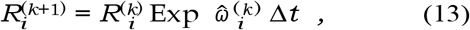

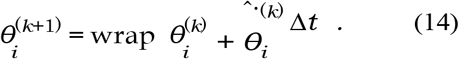

After each rotation update, 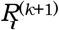 is re-orthonormalised with SVD to limit drift off SO(3). Torsions are wrapped back to the principal interval after each update. This combination preserves valid frame structure while keeping the integration step simple and efficient.

When atom-level supervision is required, the predicted frames and torsions are converted to atom14 coordinates through the OpenFold-compatible torsion-to-atom14 reconstruction pipeline. This lets the model predict in a compact residue-frame space while still receiving all-atom geometric supervision during training. It also ensures that local losses such as peptide geometry and structural violations are computed on a chemically meaningful reconstruction rather than on unconstrained raw coordinates.

The choice of a first-order integrator is deliberate for this version of the model. It keeps training and inference numerically straightforward, makes it easy to chain windows autoregressively, and avoids introducing a second learned state such as acceleration. At the same time, the experiments later show that integration error accumulation is one reason performance weakens at longer rollouts, which motivates the discussion of Velocity Verlet as future work.

### 3.5 Training Details

Training uses AdamW, BF16 mixed precision, a single NVIDIA A100, batch size 8, and a residue cap of 160. The primary objective is the velocity-space flow-matching loss; geometry losses are introduced only after this signal has stabilised. In the final schedule, FM dominates the first 150 warmup steps and heavier geometry terms ramp in after step 1000. This schedule is not incidental: without it, the combination of frame losses, atom14 supervision, and structural regularisers destabilises early optimisation. The physical-space supervision includes frame translation, frame rotation, torsion reconstruction, atom14 reconstruction, and peptide/violation-style geometry losses.

The total loss can be written schematically as

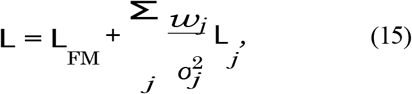

where *j* ranges over physical-space objectives and the variance terms are estimated from the training statistics of the corresponding channels. This makes the curriculum more robust across channels with different natural variances and prevents any single physical-space loss from dominating purely because of units.

About 15% of batches are converted into an endpoint-conditioned interpolation task, which asks the model to generate an internally consistent path when both start and end states are provided. This improves path consistency without changing the main task into pure interpolation. The final training setup also uses EMA decay 0.99, multi-stride supervision over {1, 2, 4}, and a three-phase global gradient clipping schedule of 500 → 50 → 15. During inference, the EMA weights are used, anchor correction is disabled (`ANCHOR_ALPHA=` 0.0), and each 16-frame window is generated with 32 Euler steps in flow time before autoregressive chaining to the next window.

## 4 Experiments

The curriculum can be summarised in three phases. Phase 1 uses FM-dominant supervision so that the network first learns a stable velocity field. Phase 2 introduces atom14 and geometry-aware losses after step 1000, when the trajectory predictor is already producing reasonable state reconstructions. Phase 3 adds the interpolation task and the later geometric penalties, which refine path consistency rather than establishing the basic motion prior from scratch. This ordering was necessary in practice: activating all losses from the start caused optimisation instability and poorer final dynamics.

### 4.1 Setup

The dataset is the filtered ATLAS MD corpus [13], yielding 710 proteins and 2090 replicate trajectories after pre-processing and protein-level splitting. We use an 80/10/10 split by protein identity so that all replicates of a protein remain in the same partition. The final training, validation, and test partitions contain 570/70/70 proteins respectively, with 1670/210/210 replicate trajectories. Trajectories are smoothed before training with a Savitzky–Golay filter (window 7, polynomial order 2), and evaluation is performed on 72 held-out proteins in the final test subset. Key implementation settings are listed in Table 1, while the overall evaluation protocol is summarized in Table 2.

**Table 1:** Key implementation settings for the final model.

| Setting | Value |
| --- | --- |
| Parameters | $\sim 15\text{M}$ |
| Single representation $C_S$ | 384 |
| Pair representation $C_Z$ | 128 |
| IPA blocks | 6 |
| Temporal layers / heads | 2 / 8 |
| Window size $W$ | 16 |
| Strides | {1, 2, 4} |
| Batch size | 8 |
| Max residues | 160 |
| EMA decay | 0.99 |
| Gradient clip | 500 → 50 → 15 |

**Table 2:** Core experimental protocol for the final evaluation.

| Item | Value |
| --- | --- |
| Proteins / trajectories | 710 / 2090 |
| Split | 80 / 10 / 10 by protein |
| Training GPU | 1× A100 |
| Window / stride | 16 / {1, 2, 4} |
| Primary rollout | 128 frames |
| Inference FM steps | 32 per window |
| Anchor correction | disabled at test time |
| Main test proteins | 72 |

The primary rollout horizon is 128 frames, corresponding to eight autoregressive windows of length *W* =16. This horizon is chosen because it matches the intended operating regime of the model and gives the best distributional agreement with ground truth: on the detailed `8htw_B` study, the top-5-PC MMD is minimised at 128 frames. We therefore treat 128 frames as the main benchmark and longer rollouts as stress tests rather than as the primary claim.

Evaluation uses three complementary metric families. *Structural accuracy*: TM-score, C*α* RMSD, and GDT-TS measure whether generated structures remain in the correct fold-recognition regime. *Geometric validity*: valid C*α*–C*α* bond rate, steric clash percentage, and Ramachandran favoured percentage assess whether the rollout remains physically plausible at residue level. *Dynamics quality*: RMSF Pearson correlation, RMSF ratio, diversity RMSD, and low-dimensional MMD measure whether the model captures the pattern and magnitude of residue flexibility and occupies the correct part of conformational space. This metric split is necessary because strong values on one family do not guarantee quality on the others.

#### Computational efficiency

Table 3 compares wall-clock inference cost against equivalent-timescale MD simulation. On a single NVIDIA A100, VelocityFM generates a 128-frame trajectory in 97 s total (59 s steady-state after a one-time 45 s `torch.compile` JIT warmup) at a peak memory cost of only 2.3 GB, yielding a **23–57**× speed-up over equivalent MD simulation. At population scale, the full 72-protein test set is evaluated in approximately 6 minutes using batched inference with 16 proteins per forward pass. These comparisons must be read alongside the accuracy caveats in Table 3: VelocityFM generates a learned approximation, not physically ground-truth dynamics.

**Table 3:** Computational cost comparison for generating a 128-frame trajectory (12.8 ns equivalent simulated time) for a single protein of about 160 residues on a single NVIDIA A100 GPU. MD cost is estimated for explicit-solvent all-atom simulation at the same timescale using OpenMM [14]. VelocityFM time is measured at inference on comparable hardware. Training cost is a one-time cost and is not included.

| Property | MD<br>(OpenMM) | VelocityFM |
| --- | --- | --- |
| Trajectory type | Physics-based | Learned approximation |
| 128-frame wall time | 92 min | 97 s (1.6 min) |
| Steady-state per protein | 92 min | 59 s (after warmup) |
| Peak GPU memory | 16 GB | **2.3 GB** |
| Single-protein speed-up | 1× | 23–57× |
| 72-protein batch time | >44 hr | **about 6 min** |
| Force field required | Yes | No |
| Scales to new protein | Full re-run | Single forward pass |
| Structural accuracy | Ground truth | TM-score<br>0.929 (median) |

<sup>a</sup> Estimated from 200–500 ns/day for an approximately 160-residue protein in explicit solvent on an A100 GPU [14].
<sup>b</sup> Includes a one-time 45 s `torch.compile` JIT warmup; steady-state inference time is 59 s.
MD and VelocityFM do not produce identical outputs: MD generates physics-based trajectories, whereas VelocityFM generates a learned approximation trained on MD data.

### Code and Data Availability

All materials produced in this research are publicly accessible via the shared resource folder at: https://drive.google.com/drive/folders/1vrR18YjzwUs5Sz-MOtK280VMmGTn52ux

The folder is organised into five components. (1) The model training notebook (Google Colab) implements the full VelocityFM training pipeline, including the data loader, loss curriculum, and checkpoint management: https://colab.research.google.com/drive/1r7dt69QwYjYwBffr8fpwTBIQzFZsE-MD

(2) The inference notebook (Google Colab) generates 128-frame trajectories from any PDB input and reproduces the 72-protein evaluation reported in Section **??**: https://colab.research.google.com/drive/1NWolICVYZXOHvnpmx5SGolGkiojVJPA3

(3) Additional analysis notebooks covering the validation pipeline, population-level metric computation, and figure generation: https://drive.google.com/drive/folders/1sN4fy4MRC_Bk_OinLmutUSbS3my0hZ_e

(4) Processed ATLAS dataset splits containing NPZ files for all 710 filtered proteins in training, validation, and test partitions: https://drive.google.com/drive/folders/1nBC8Ak3GsMzOBh2D_AHcaua6-W1uK300

(5) Training runs folder containing the final checkpoint from Training Run 192, JSONL training logs, and intermediate results: https://drive.google.com/drive/folders/1l7aoHdzbhyoQQbYx91qKNWq7QYsR6Ez3

All notebooks run on Google Colab and include Drive mount instructions.

### 4.2 Multi-Frame Degradation Analysis

Table 4 evaluates `8htw_B` from 8 to 512 frames. The main pattern is graceful degradation rather than collapse. TM-score remains above 0.92 through 128 frames, C*α* RMSD rises only from 1.45 Å at 16 frames to 1.59 Å at 128 frames, and all generated rollouts remain clash-free. This already shows that the learned rollout is stable over multiple autoregressive windows.

**Table 4:** Multi-frame degradation analysis for 8htw_B. Ground-truth values are shown for reference.

| Metric | GT | 8f | 16f | 32f | 64f | 128f | 256f | 512f |
| --- | --- | --- | --- | --- | --- | --- | --- | --- |
| TM-score | 0.955 | 0.939 | 0.941 | 0.939 | 0.931 | 0.921 | 0.890 | 0.809 |
| Cα RMSD (Å) | 1.21 | 1.49 | 1.45 | 1.46 | 1.52 | 1.59 | 1.88 | 2.64 |
| Helix % | 42.7 | 41.7 | 41.3 | 41.8 | 38.9 | 31.6 | 24.8 | 19.2 |
| Rama fav. % | 77.2 | 80.5 | 80.0 | 79.7 | 79.8 | 79.5 | 77.2 | 73.4 |
| RMSF ratio | 1.000 | 0.258 | 0.321 | 0.427 | 0.533 | 0.669 | 0.998 | 1.617 |
| RMSF Pearson $r$ | 1.000 | 0.512 | 0.732 | 0.618 | 0.616 | 0.611 | 0.543 | 0.387 |
| Valid bond % | 86.9 | 75.4 | 80.9 | 79.2 | 73.3 | 72.1 | 73.6 | 71.3 |
| Clash-free % | 100 | 100 | 100 | 100 | 100 | 100 | 100 | 100 |
| MMD (5 PCs) | 0.000 | 11.33 | 10.78 | 10.03 | 9.19 | 8.94 | 14.35 | 26.22 |

The table also clarifies the structure–dynamics trade-off. At short horizons, secondary structure and Ramachandran quality are very strong: helix content is 41.3% at 16 frames versus 42.7% in ground truth, and Ramachandran favoured percentages remain near or above the reference level through 128 frames. Motion amplitude, however, is initially under-dispersed, as shown by the RMSF ratio rising from 0.258 at 8 frames to 0.669 at 128 frames.

The key result is that 128 frames gives the best overall operating point. Its top-5-PC MMD is the minimum across the tested horizons, indicating the best match to the reference conformational distribution. At 256 frames the RMSF ratio becomes nearly perfect (0.998), but this comes with weaker structure preservation and much worse MMD. In other words, 256 frames calibrates amplitude 6better, whereas 128 frames give the best combined structural and distributional behaviour.

This behaviour is exactly what one would expect from a model trained on short windows and rolled out autoregressively. The early horizons are structurally easiest because little error has accumulated, whereas longer horizons allow amplitude to grow but also amplify integration and prediction drift. The 128-frame point is therefore not chosen because it gives the best single metric, but because it gives the best overall balance across structure, geometry, and conformational distribution.

### 4.3 Population-Level Results

Table 5 summarises the 72-protein benchmark at 128 frames. The strongest finding is complete fold-recognition generalisation: all 72 proteins remain above TM*>* 0.7. Structural quality is high overall, with median TM-score 0.929 [0.909, 0.941], median C*α* RMSD 1.343 [1.138, 1.526] Å, and median GDT-TS 0.846 [0.825, 0.883]. These are strong values for a learned short-horizon rollout model that predicts on unseen proteins rather than memorising individual trajectories.

**Table 5:** Population-level results on 72 held-out proteins at 128 frames. Values are median [Q25, Q75].

| Metric | Median [Q25, Q75] |
| --- | --- |
| TM-score | 0.929 [0.909, 0.941] |
| % frames TM > 0.7 | 100% [100%, 100%] |
| Cα RMSD (Å) | 1.343 [1.138, 1.526] |
| GDT-TS | 0.846 [0.825, 0.883] |
| Valid Cα-Cα % | 72.96% [70.50%, 77.07%] |
| Steric clash % | 0.000% [0.000%, 0.000%] |
| Ramachandran fav. % | 91.09% [89.48%, 93.01%] |
| RMSF Pearson $r$ | 0.663 [0.571, 0.772] |
| RMSF ratio | 0.697 [0.552, 0.917] |
| Helix preservation ratio | 0.737 [0.690, 0.769] |
| Diversity RMSD (Å) | 0.625 [0.560, 0.725] |

Geometry also generalises strongly. All proteins are clash-free, valid C*α*–C*α* percentage is 72.96% [70.50%, 77.07%], and the median Ramachandran favoured rate is 91.09% [89.48%, 93.01%]. This means the model is not merely preserving coarse fold topology; it is also maintaining valid residue-level backbone geometry across the held-out set. Importantly, the zero-clash result holds for every protein rather than only in aggregate, indicating that the learned self-avoidance prior is highly stable under rollout.

The remaining weakness appears in flexibility statistics rather than geometry. Median RMSF Pearson correlation is 0.663 [0.571, 0.772], which indicates that the residue-wise pattern of motion is often moderately aligned with the reference trajectory. The RMSF ratio, however, is 0.697 [0.552, 0.917], indicating that the predicted motion amplitude is systematically smaller than ground truth. This distinction between pattern and amplitude is central to interpreting the model correctly.

Helix preservation ratio adds another useful perspective. Its median value of 0.737 [0.690, 0.769] means that the model retains most, though not all, native helical content at the main horizon. This is much stronger than the much longer-horizon failure cases seen in earlier exploratory rollouts and aligns with the claim that the present model should be judged in the short-horizon regime for which it was designed.

Figure 2 makes the central population result visually clear. The model is consistently inside the fold-recognition regime on every test protein. This matters because it converts the evaluation from a mean-performance claim into a generalisation claim: the model is not succeeding only on easy proteins or on a small favourable subset. The same plot also shows that proteins with stronger ground-truth structural stability tend to remain easier for the generator, but the overall spread is narrow enough that fold preservation is not restricted to one narrow class of targets.

**Figure 2:**
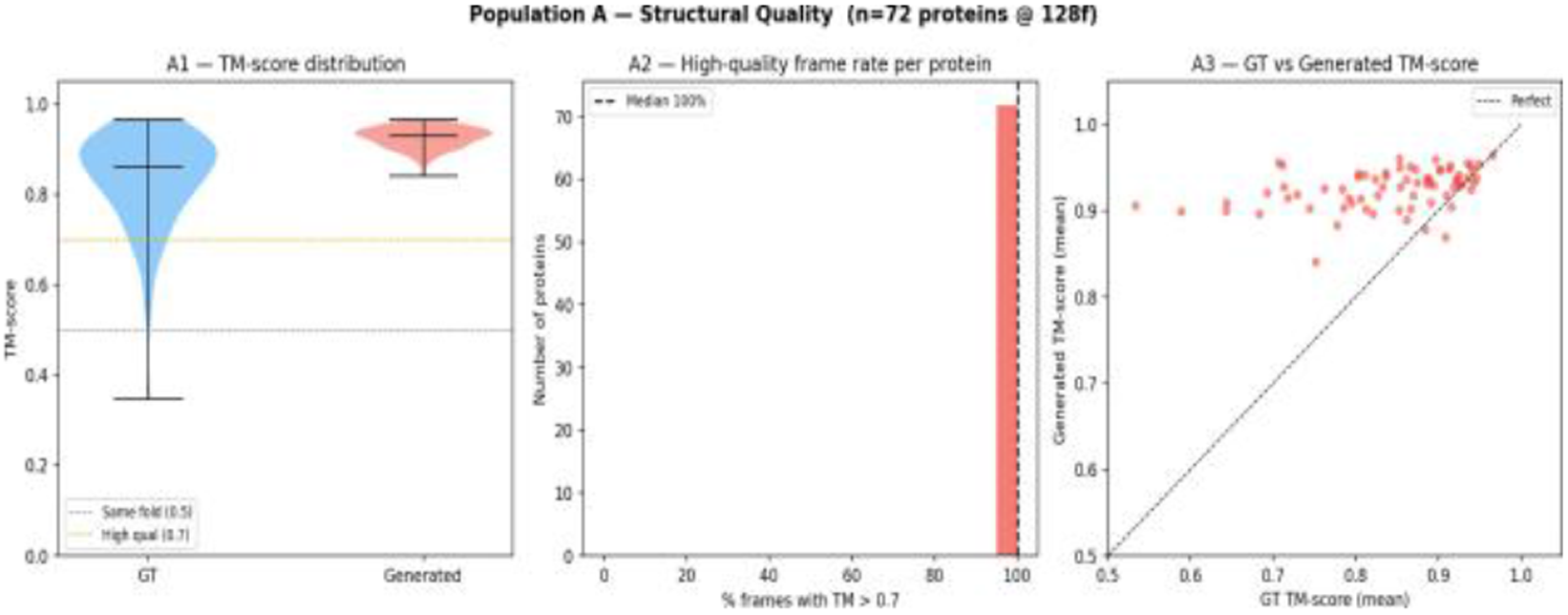
Population-level structural quality across 72 held-out test proteins at 128-frame rollout. All 72 proteins maintain TM-score *>* 0.7 (centre panel, 100% bar).

### 4.4 Case Studies

The aggregate statistics hide an important qualitative spread, so we examine four held-out proteins in more detail. Representative examples are shown in Figure 3. Each case preserves the broad fold, but the agreement in flexibility statistics varies considerably. This makes the case studies useful not only as examples of success and failure, but also as evidence that the task decomposes into at least two partially separate subproblems: staying in the correct structural basin and calibrating the local motion pattern within that basin.

**Figure 3:**
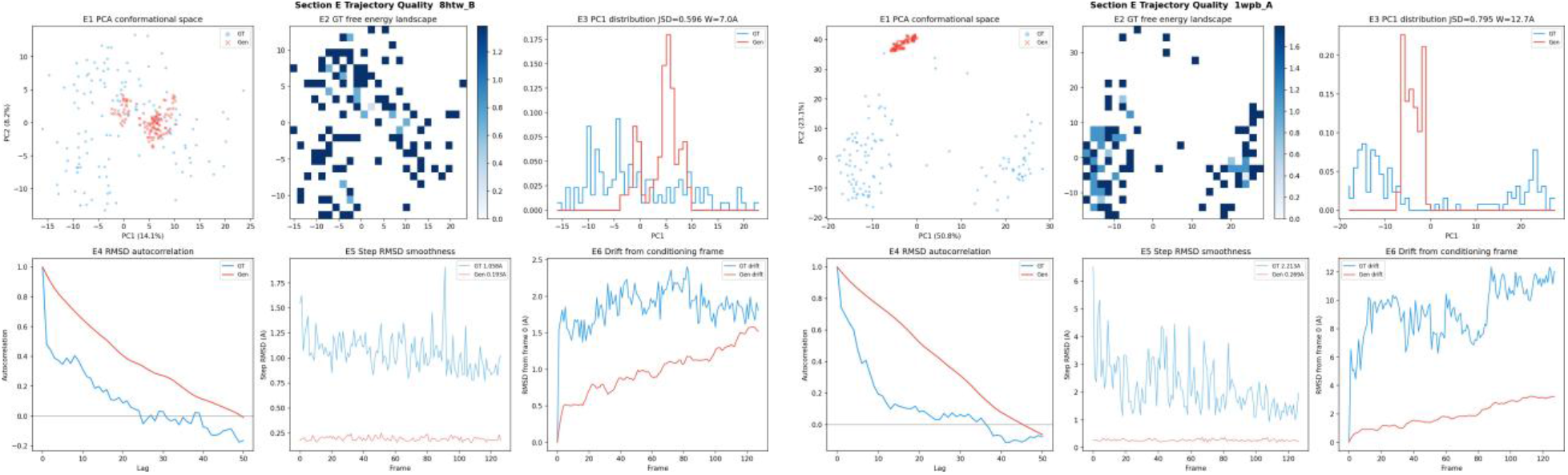
Protein-specific trajectory case studies. Left: `8htw_B,` an easier helical target where short-horizon rollout remains compact and structurally stable. Right: `1wpb_A,` a more dynamic target where the fold is still preserved but the motion becomes more conservative and harder to calibrate. These examples visually reinforce the quantitative gap between fold preservation and dynamics fidelity.

`8htw_B` is a helical 181-residue target and represents the intended operating regime of the model. It achieves TM= 0.941 and RMSF *r* = 0.732, while the multi-frame analysis shows only gradual degradation up to 128 frames. Qualitatively, the trajectory remains compact and helices are preserved across the rollout, which is consistent with the low RMSD and stable geometric-validity metrics. This case therefore supports the claim that the residue-frame representation and velocity-space objective are well matched to short-horizon, locally smooth conformational motion.

`1qw2_A` is the best dynamics case in the held-out set, with TM= 0.925 and RMSF *r* = 0.921. Here the model does not merely preserve the fold; it also reproduces the per-residue flexibility ranking with high fidelity. This is an important positive control because it shows that the conservative RMSF ratio seen in the population statistics does not imply a fundamental inability to model dynamics. Rather, the architecture can recover both structure and motion when the target trajectory remains close to the dominant basin learned during training.

`1lfp_A` is the most informative decoupling case. Its TM-score is very high at 0.944, yet RMSF correlation falls to 0.270. Figure 4 shows that the generated ensemble remains in the correct structural region while failing to reproduce the internal spread of the reference distribution. In other words, the model knows *which fold* to stay in, but not *how motion should be distributed* inside that fold. This distinction is scientifically important because a structure-only evaluation would classify the example as a clear success, whereas a dynamics-aware evaluation reveals a real limitation.

**Figure 4:**
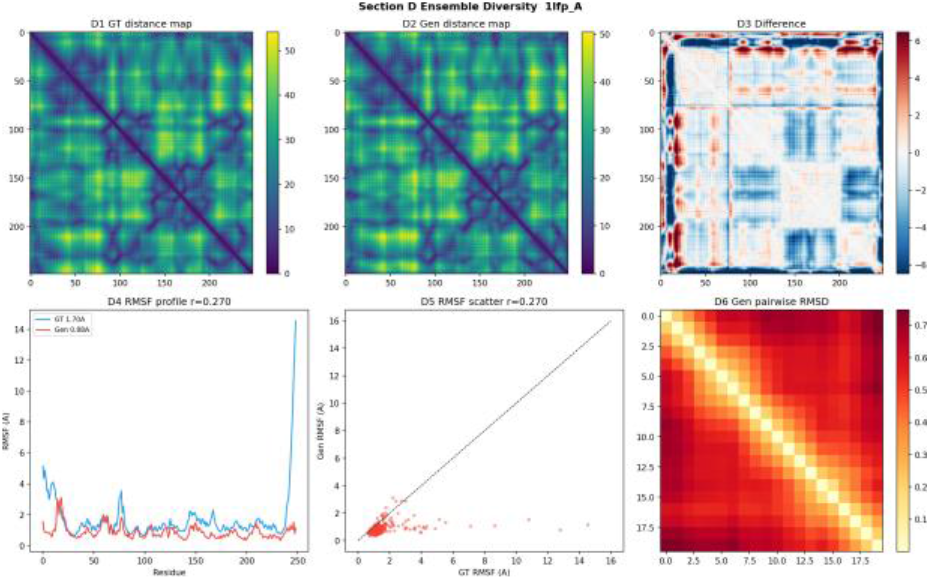
Case-study visual for `1lfp_A,` the fold–dynamics decoupling example. The generated ensemble stays in the correct structural basin but undercaptures the internal diversity pattern of the reference trajectory.

`1wpb_A` is the most dynamic target among the four, with ground-truth diversity 5.15 Å, TM= 0.882, and RMSF *r* = 0.323. It remains recognisably folded, but the rollout becomes visibly more conservative and less well calibrated than in `8htw_B or 1qw2_A`. This is the clearest indication that large-amplitude motion is still the hardest regime for VelocityFM. Taken together, the four cases suggest that fold preservation generalises relatively uniformly, whereas flexibility calibration depends much more strongly on the dynamical complexity of the target protein.

### 4.5 RMSF Calibration

Figure 5 summarizes flexibility calibration across the population. The median RMSF Pearson correlation of 0.663 shows that the model captures a meaningful residue-wise pattern of flexibility, but the median RMSF ratio of 0.697 shows that the amplitude is conservative at 128 frames. The architecture therefore knows *where* motion tends to occur better than *how much* motion should occur.

**Figure 5:**
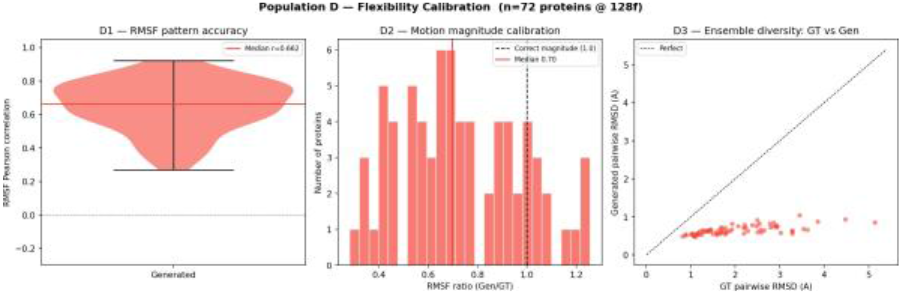
Flexibility calibration across 72 proteins. RMSF Pearson *r* = 0.663 (median) shows moderate per-residue dynamics agreement. RMSF ratio 0.697 indicates conservative motion generation at 128 frames, consistent with the single-protein analysis.

This interpretation is consistent with both the population and the single-protein analyses. On `8htw_B,` RMSF correlation is reasonable at 128 frames, but amplitude remains below the ground-truth level. On the harder cases, especially `1lfp_A` and `1wpb_A,` the mismatch becomes more obvious. These patterns explain why VelocityFM performs very strongly on structure-preserving metrics while remaining conservative as a motion generator. They also show why RMSF calibration must be reported alongside TM-score: a model can look excellent structurally while still failing to reproduce the correct dynamical scale.

## 5 Discussion

VelocityFM should be interpreted as a strong *short-horizon fold-preserving predictor*, not as a replacement for long-horizon MD. At the intended 128-frame horizon, the evidence is consistent across metrics: median TM-score is 0.929, every held-out protein remains above TM*>* 0.7, all rollouts are clash-free, and backbone geometry remains valid with 91.09% median Ramachandran favoured rate. The architecture therefore succeeds on the core CS/ML objective of learning stable local trajectory evolution on unseen proteins.

The main limitation is motion calibration rather than structural correctness. The median RMSF ratio of 0.697 shows that the generator is conservative, and the `1lfp_A` case shows that fold quality and dynamics quality can decouple sharply even when the backbone remains in the correct basin. Two experimental limitations also matter. First, the split is by protein identity rather than by a strict sequence-homology threshold, so future evaluation should be more stringent. Second, the present study is limited to single-chain proteins and short autoregressive horizons; performance degrades beyond the main operating regime. These caveats mean the current model should be read as a trajectory predictor within a bounded horizon, not as a full MD surrogate.

The next improvements are technically clear. Velocity Verlet integration could reduce integration error accumulation during autoregressive rollout. A stronger spatial block such as an AtomTransformer-style encoder could improve local geometry and motion distribution modelling. A multi-step rollout loss could explicitly penalise error accumulation across chained windows rather than supervising each window mostly in isolation. Together, these changes target exactly the failure modes diagnosed in the experiments: cumulative integration drift, conservative motion amplitude, and the remaining gap between fold fidelity and dynamics fidelity.

## 6 Conclusion

We presented VelocityFM, a residue-frame trajectory model that applies rectified flow matching in velocity space to short-horizon protein dynamics prediction. The final system combines geometric spatial reasoning, temporal attention, and staged physical supervision, and generalises to 72 unseen proteins with median TM-score 0.929, 100% TM*>* 0.7, and 100% clash-free generation at 128 frames. Its remaining weakness is conservative motion amplitude rather than structural breakdown, making it a useful short-horizon predictor but not a substitute for unrestricted MD simulation. Extending the model toward better integration schemes, stronger spatial encoders, and rollout-aware objectives is a concrete path toward more realistic learned protein dynamics.

